# Herbarium records reveal early flowering in response to warming in the southern hemisphere

**DOI:** 10.1101/432765

**Authors:** Barnabas H. Daru, Matthew M. Kling, Emily K. Meineke, Abraham E. van Wyk

## Abstract

**Premise of the Study:** Herbarium specimens are increasingly used as records of plant flowering phenology, which has advanced for many species in response to climate change. However, most herbarium-based studies on plant phenology focus on taxa from temperate parts of the northern hemisphere. Here, we explore flowering phenologic responses to climate in a temperate/subtropical plant genus *Protea* (Proteaceae), an iconic group of woody plants with year-round flowering phenology and endemic to sub-Saharan Africa. *Protea* is widely used in horticulture and is a flagship genus for the flora of the hyperdiverse Cape Floristic Region.

**Methods:** We used a database of 2154 herbarium records of 25 *Protea* species to explore patterns in flowering spanning the past 100 years. We used a circular sliding window analysis to characterize phenological patterns in these aseasonal species, plus a novel linear mixed effects model formulation to test how both site-to-site and year-to-year variation in temperature and precipitation affect flowering date across species.

**Results:** Both warmer sites and warmer years were associated with earlier flowering of 3–5 days/°C. In general, the timing of peak flowering was influenced more strongly by temperature than precipitation. Although species vary widely in when they flower during the year, their phenological responses to temperature are phylogenetically conserved, with closely related species tending to shift flowering time similarly with increasing temperature.

**Discussion:** Together, our results point to climate-responsive phenology for this important plant genus. Our results indicate that the subtropical, aseasonally-flowering genus *Protea* has temperature-driven flowering phenologic responses that are remarkably similar in magnitude to those of better-studied northern temperate plant species, suggesting a generality across biomes that has not been described elsewhere.

## Introduction

Phenology is the study of periodic life cycle stages, especially how these are influenced by weather and climate (Schwartz, 2013). Phenological change in response to climate change is a topic of growing interest in ecology as well as conservation and evolutionary biology. In particular, changes in flowering phenology are likely to have downstream effects on important ecosystem processes and biotic interactions, including impacts on the various animal taxa that depend on plants for pollen, nectar, or fruit/seed (Fitter and Fitter, 2002; Parmesan and Yohe, 2003; Visser and Both, 2005; Post and Inouye, 2008).

Whereas some plant species have demonstrated trends toward earlier phenological events with anthropogenic climate change (Hart et al., 2014), others have shown either a delay in the onset of flowering in response to warming (Fitter and Fitter, 2002) or no significant change (Miller-Rushing and Primack, 2008). In part, this is thought to be because different plant species use different cues to time phenological events. For example, in many plants, day length (photoperiod) has traditionally been considered more important than low winter temperature (vernalization) in synchronising flowering to the changing seasons (e.g., Yanovsky and Kay, 2003; Saikkonen et al., 2012), but an interaction between both (and other) cues has been demonstrated in *Arabidopsis thaliana* (Andrés and Coupland, 2012).

Perhaps as a result of related species using similar cues, phenological shifts are seemingly non-random across lineages (Willis et al., 2008; Davis et al., 2010; Davies et al., 2013), emphasizing the need to explore phenological change within a phylogenetic framework. However, the phylogenetic conservatism of phenological response has only been tested on a small subset of species (Davies et al., 2013), and has not been explored for entire plant communities with fine-scale phylogenetic resolution, nor across the broad distributional ranges of numerous co-occurring species. If phenological responsiveness to climate is phylogenetically patterned within communities—within or between lineages—phylogenetic information may have value for understanding phenological cuing mechanisms and forecasting future responses to climate change.

However, phenology data for assessing patterns and processes in phenological change are sparse. Long-term observational data on flowering, leaf-out, and fruiting are limited across space, time, and clades, and short-term warming experiments do not reliably reproduce the effects of long-term climate change (Wolkovich et al., 2012). A critical bias in long-term phenology data is that they are available primarily for temperate regions and only in rare cases for the tropics, where most plant diversity occurs. One potential way to overcome the constraints of long-term field observational data on phenophases is by using historical records in herbaria and museums (Davis et al., 2015). Although such records have not necessarily been collected expressly for phenological investigations, and therefore present their own biases (Daru et al., 2018), a significant body of literature now exists in which historical records have potential for investigating climate-related phenological trends across plant species (Primack et al., 2004; Bolmgren and Lonnberg, 2005; Coleman and Brawley, 2005; Lavoie and Lachance, 2006; Miller-Rushing et al., 2006; Bowers, 2007; Houle, 2007; Kauserud et al., 2008; Gallagher et al., 2009; Neil et al., 2010; Park and Mazer, 2018). Like observational studies, however, most herbarium-based studies on plant phenology are established on collections from temperate parts of the northern hemisphere. Collections remain almost completely untapped for determining phenological responses to climate change in the tropics and subtropics, as well as temperate parts of the southern hemisphere.

Here, we investigate phenological change for *Protea* (Proteaceae), commonly known as proteas or sugarbushes, an iconic flowering plant genus endemic to sub-Saharan Africa, with its center of diversity in southern African (Rourke, 1982). *Protea* is comprised of woody shrublets, shrubs or trees displaying a suite of floral adaptations (Figure 1) for a diverse array of pollinators including beetles, birds and small mammals (Collins and Rebelo, 1987; Wright et al., 1991). In southern Africa, flowering in most species of *Protea* is hypothesized to peak in spring and summer, with only a few species in autumn, perhaps reflecting the abundance of pollinators (Rebelo, 2001). In a phenology-based study involving four species of *Protea* confined to the core Cape Floristic Region, it was shown that arthropod communities are significantly influenced by infructescence phenology (Roets et al., 2006). *Protea* is widely used in horticulture (mainly as cut flowers) and is a flagship genus for the flora of the Cape Floristic Region/Kingdom (Vogts and Patterson-Jones, 1982; Gollnow and Gerber, 2015).

**Figure 1.**
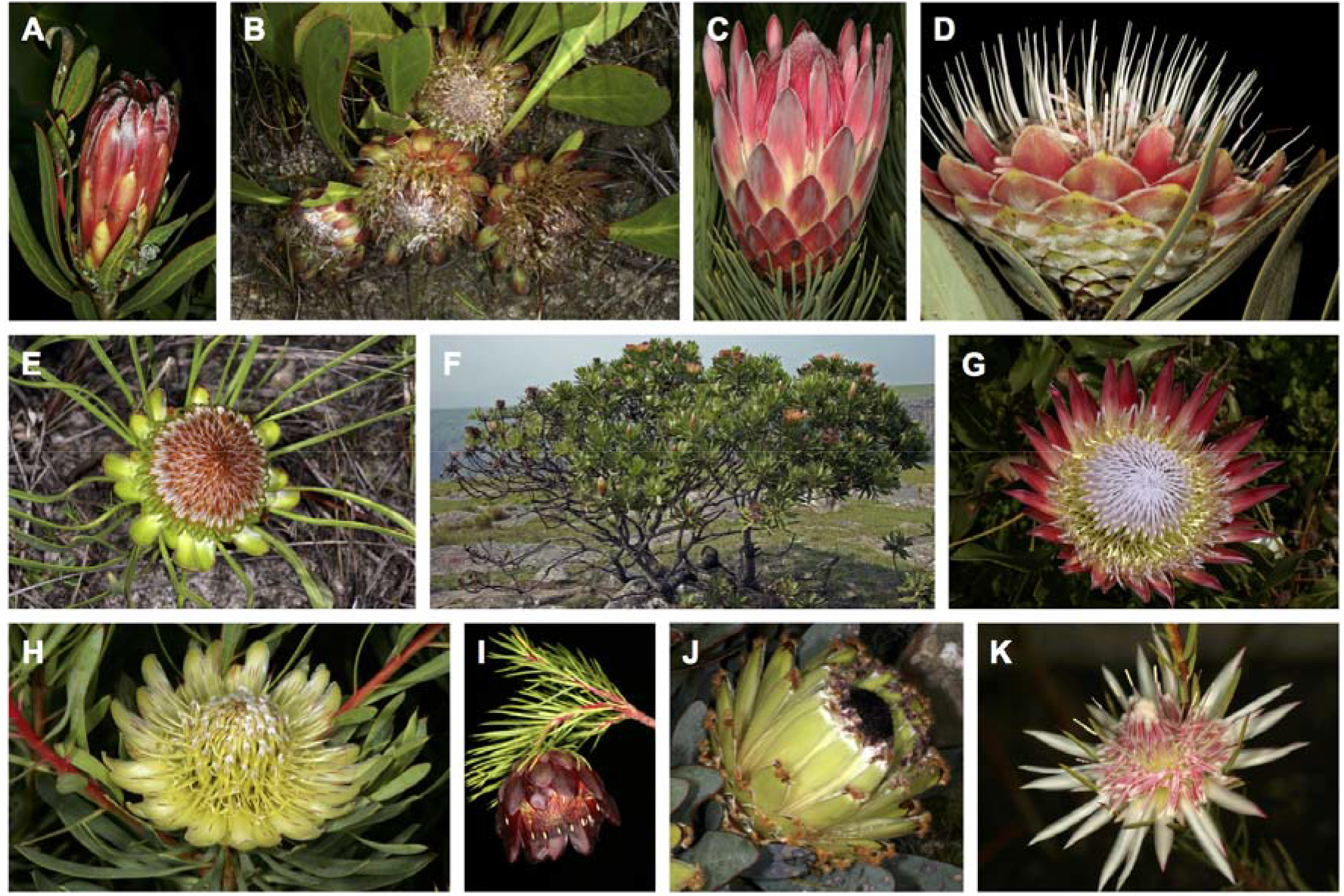
Representatives of the ∼115 species of *Protea* (Proteaceae) showing variation in flowerhead and growth form. (A) Flowerhead of *Protea burchellii*, Caledon, South Africa; (B) *P. acaulos*, Sir Lowry’s Pass, Somerset West; (C) *P. aristata* flowerhead; (D) *P. caffra* subsp. *caffra*, Fairy Glen Nature Reserve, Pretoria; (E) *P. angustata* Kleinmond; (F) Green-leaved form of *P. roupelliae* subsp. *roupelliae* tree in flower, Mtentu River Gorge, Eastern Cape; (G) flowerhead of *P. cynaroides*; (H) *P. scolymocephala*; (I) *P. nana*, Paarlberg Nature Reserve, Paarl; (J) *P. magnifica*; and (K) *P.mucronifolia*. Pictures A–K courtesy of SAplants [CC BY-SA 4.0], Wikimedia Commons.

The genus *Protea* comprises about 115 species, the bulk (± 70) of which are endemic or near-endemic to the Cape Floristic Region (Manning and Goldblatt, 2012).

Climatically the Cape Floristic Region is temperate (Van Wyk and Smith, 2001), with mean annual temperatures varying from 15–16 °C at the coast to 17–18 °C in interior areas, but are lower than 13 °C at higher altitudes. Frost is restricted to the inland valleys, and snow falls on the higher mountains. The western part of the Cape Floristic Region receives most of its rainfall in winter (so-called Mediterranean climate), but to the east the rainfall is more evenly distributed throughout the year. Average annual rainfall is mostly between 300 and 2000 mm, but it is estimated to be as high as 5000 mm on some of the mountain peaks. Elsewhere in southern Africa the geographic range of *Protea* falls mainly in areas with summer rainfall (± 14 species), while about 35 species are dispersed further north in central and tropical Africa (Rourke, 1982). *Protea* is clearly a temperate to subtropical floristic element, and outside the Cape it is usually associated with either more temperate (compared with surrounding subtropical and tropical conditions) high-altitude habitats or the cooler south-facing aspects of hills and mountains. In sub-Saharan Africa these more temperate climatic conditions are one of the defining features of the Afromontane and Afroalpine Archipelago-like Centre of Endemism (White, 1983; Huxley et al., 1998), a biogeographical region with floristic links to Cape Floristic Region. Hence *Protea* may also be considered an Afromontane floristic element.

Despite its considerable ecological significance and cultural importance, the flowering phenology of *Protea* remains poorly studied (Pierce, 1984; Johnson, 1993). Even in the case of commercially grown plants, the factors that trigger flowering onset are poorly understood. The great variation that exists in *Protea* flowering times and apparent flowering prerequisites, suggests multi-factorial control of the *Protea* flowering cycle (Hoffman, 2006). In particular, for some cultivated members of *Protea*, e.g. *Protea cynaroides* (Figure 1), flowering may not depend strictly on photoperiod, and a cold treatment during winter is apparently required by some species to trigger flowering (Hoffman, 2006, and references therein), whereas in the case of resprouters, the age of vegetative growth since the last fire may also play a role (Rebelo, 2008).

In this study, we use a database of 7770 herbarium specimens collected between 1808 and 2011 to explore the potential for phenological shifts in flowering across the genus *Protea*. Specifically, we: (i) characterize seasonal and geographic flowering phenology patterns across *Protea* species, (ii) investigate how site-to-site and year-to-year variation in temperature and precipitation influence *Protea* flowering phenology, and (iii) test for phylogenetic conservatism in these climatic effects on phenology. Our study reveals how an iconic plant genus in an understudied part of the world responds to climate variation across space and time, and we discuss how this might cascade to affect its native ecosystem.

## Materials and Methods

### Herbarium data

We compiled information from 7770 herbarium specimens for 87 species of *Protea* (Proteaceae). The data were collected from herbaria across South Africa as archived at the National Herbarium of the South African National Biodiversity Institute in Pretoria, and include records from Swaziland and Lesotho as well as South Africa. From each of these specimens we extracted four types of data: species identity, collection date (year, month, day), geographic coordinates (often represented as quarter degree grid cells, QDGC system) and flowering status (whether or not the specimen was in peak flower). Each QDGC was converted to decimal degrees following Larsen et al. (2009). For example a QDGC of 3419AD = longitude 19.375 and latitude -34.375; or 3318CD = longitude 18.375 and latitude -33.875, and so on (Larsen et al., 2009). Peak flowering was assessed by examining each specimen and determining whether more than 50% of flower buds (of at least one open flowerhead) were in anthesis, a technique that has been used in other studies to represent flowering phenology (Primack et al., 2004; Hart et al., 2014). Specimens not in peak flower were removed from the analysis. We also removed records falling farther north than the northernmost point in South Africa, records from before 1900, and records listed as collected on the last day of any month (these dates were dramatically overrepresented in the data, suggesting that they were often used as a default when the true collection day was unknown). Lastly, species with fewer than 50 remaining records were removed. After data cleaning, the final dataset included 2154 records representing 25 species of *Protea*.

### Phenological patterns

For each species, we used the frequency distribution of specimen collection dates as a proxy for flowering phenology (Panchen et al., 2012). We converted dates on specimens to Julian Day of Year (DOY; where January 1 = 1 DOY and February 1 = 32 DOY, and so on). To characterize the flowering phenology for each species, we defined the “peak season” as the center of the sliding three-month window with the largest number of occurrences, “dormant season” as the center of the sliding six-month window with the fewest number of occurrences, and “aseasonality” as the ratio of the number of dormant season occurrences to the number of peak season occurrences. These sliding windows were wrapped to reflect the circular nature of the calendar year, in which December 31 is adjacent to January 1. For each species, we calculated an adjusted version of the Julian DOY of collection for each herbarium specimen, day of flowering year (DOFY), measured as the number of days since the middle of a species’ dormant season, to allow DOY to be treated as a linear variable with no disjunction between December 31 and January 1.

To test the assumption that the seasonal distribution of herbarium specimens with 50% open flowers is a good representation of actual peak flowering in the field, we compared the center of each species’ peak flowering season derived from our dataset to phenograms provided in the “*Protea* Atlas” of Rebelo (2001), which represents the most comprehensive treatment of all described *Protea* species, including relevant information on the ecology, spatial distributions, and species abundance.

### Climate analyses

Studies have shown that the distribution of plant species richness in southern Africa is driven largely by shifts in rainfall and temperature regimes (O’Brien, 1993; O’Brien *et al*., 1998, 2000). We used temperature and precipitation data from the University of Delaware Air Temperature & Precipitation dataset version 3.01 (Willmott and Matsuura, 2001), which includes 0.5-degree gridded monthly mean temperature and precipitation for every month of every year from 1950–2010. We extracted the full climate time series (2 variables × 12 months × 61 years) at the collection location of every herbarium specimen. Precipitation values were log-transformed for normality and for ecological realism.

We used these data to explore the roles of spatial and temporal climate variation in driving *Protea* flowering dates. For each specimen record, a spatial and a temporal climate anomaly were calculated for temperature and precipitation, for a total of four predictor variables. Spatial anomalies were calculated as the difference between a location’s long-term mean climate (across all months and years in the climate dataset) and the species-wide average long-term mean climate across all specimen locations, with positive values representing specimens from warmer or wetter parts of a species range and negative values representing specimens from cooler or drier parts of the range. Temporal anomalies were calculated as the difference between annual temperature or precipitation at the collection location in the year the specimen was collected and the long-term mean climate at that location, with positive values representing collections in years that were locally warmer or wetter than average and negative values in years that were locally colder or drier than average. Since species flower at different times of year, these annual temperature anomalies were defined differently for each species, calculated as the average across the 12 month period from 8 months before through 3 months after a species’ peak flowering month; this asymmetrical window was chosen in order to encompass climate during a given peak flowering season and the preceding dormant season, which together are likely to influence flowering phenology. Normalized specimen collection date, the dependent variable, was calculated as the difference between each specimen’s DOFY as described above and the average DOFY for that species.

We fit a single mixed effects model using the R package lme4 (Bates et al., 2015), predicting normalized specimen collection date as a function of these four climate variables (spatial temperature anomaly, spatial precipitation anomaly, temporal temperature anomaly, and temporal precipitation anomaly), with random effects of species on slopes but not intercepts (since intercepts are by definition zero as a result of the de-meaning described above). This hierarchical modeling approach allows the simultaneous estimation of each climatic predictor on *Protea* flowering phenology both overall and for each individual species, both levels of which are of interest in this study. The maximum likelihood optimization criterion was used over restricted maximum likelihood, to allow model comparison. To test for the significance of each of the predictors, the full model was compared to four reduced models, each with one of the predictors removed.

### Phylogenetic analysis

Phylogenetic relationship among *Protea* species was reconstructed using DNA sequences from four plastid (*trnL, trnL-trnF, rps16* and *atpB-rbcL*) and two nuclear regions (ITS and *ncpGS*) available from GenBank. The sequences were aligned using seaview v.4.5.4 (Gouy et al., 2010) and manually adjusted using mesquite v.2.5 (Maddison and Maddison, 2008). The combined data set comprised 3386 nucleotides in length.

In a second step, we reconstructed phylogenetic relationships on the combined dataset using maximum likelihood (Stamatakis et al., 2008) via the CIPRES gateway (Miller et al., 2009). Branch lengths were transformed to millions of years by enforcing topological constraints assuming the APG III backbone from Phylomatic v.3 (Webb and Donoghue, 2005). We then used BEAUti v.1.7.5 (Drummond and Rambaut, 2007) to generate an XML file as input file for reconstructing the dated phylogenetic tree using Bayesian inference and one independent fossil calibration with normal prior distribution as follows: *Protea* root node (28.4 Ma, SD 2 Ma). We carried out a Bayesian analysis by running four chains simultaneously for 2 million generations and discarding the first 20 per cent of trees as burn-in. The distribution of posterior probabilities from the different chains were assessed by constructing a 50% majority rule consensus tree for further analysis.

We then used the dated phylogenetic tree to estimate phylogenetic signal in phenologies: whether closely related species tend to exhibit similar phenologies or diverge in the timing of reproductive events. We used Abouheif’s C_mean_ statistic (Abouheif, 1999), Blomberg’s K (Blomberg et al., 2003), and Pagel’s lambda (λ) (Pagel, 1999). Significance was assessed by shuffling the trait values 1000 times across the tips of the phylogeny, and comparing it to expectations under a Brownian or by random models. Values of Blomberg’s K and Pagel’s λ > 1 indicate high phylogenetic signal, i.e. closely related species share similar traits than expected by chance. Both Blomberg’s K and Pagel’s λ were calculated using the R package phytools (Revell, 2012), whereas Abouheif’s C_mean_ was calculated using adephylo (Jombart and Dray, 2008).

## Results

### Peak flowering over time and across geographic space

We found wide variation in the distribution of flowering phenology across geographic space, climate, and time. Temporally, species are not evenly distributed through the course of time, with the early years showing sparse records, and high density of collecting between 1960s and 1980s (Figure 2). We found a strong correlation between collection day of the year from herbarium specimens and flowering time as recorded in the literature (Rebelo, 2001; r = 0.93, Figure S1), supporting our claim that collection dates on herbarium specimens can serve as surrogate for flowering time in *Protea*. While most species tend to flower throughout the year, each species shows a distinct peak of flowering phenology (Figure S2).

**Figure 2.**
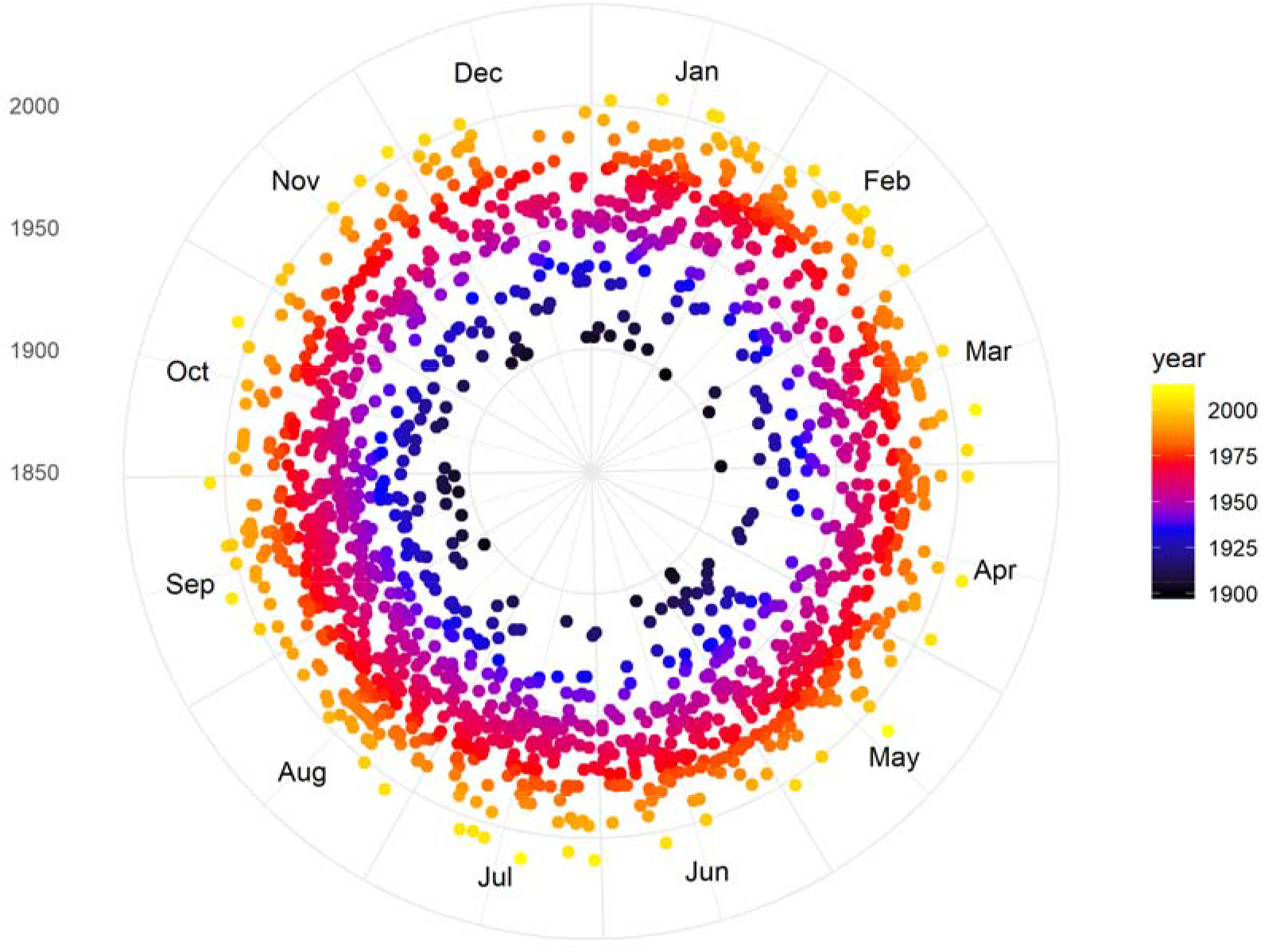
Temporal distribution of collection date of all herbarium specimens of *Protea* species in South Africa. Distance from the center represents collection year, with the inner rings indicating the early years of collecting, whereas the outer rings correspond to more modern collecting.

Geographically, we found a spatial gradient in peak flowering season across *Protea* species (Figure 3). In the Mediterranean-type Cape Floristic Region, flowering time for most species tended to peak in the winter, whereas the non-Mediterranean regions extending from Mtata in the Eastern Cape Province to Limpopo in the north show more flowering during summer (Figure 3).

**Figure 3.**
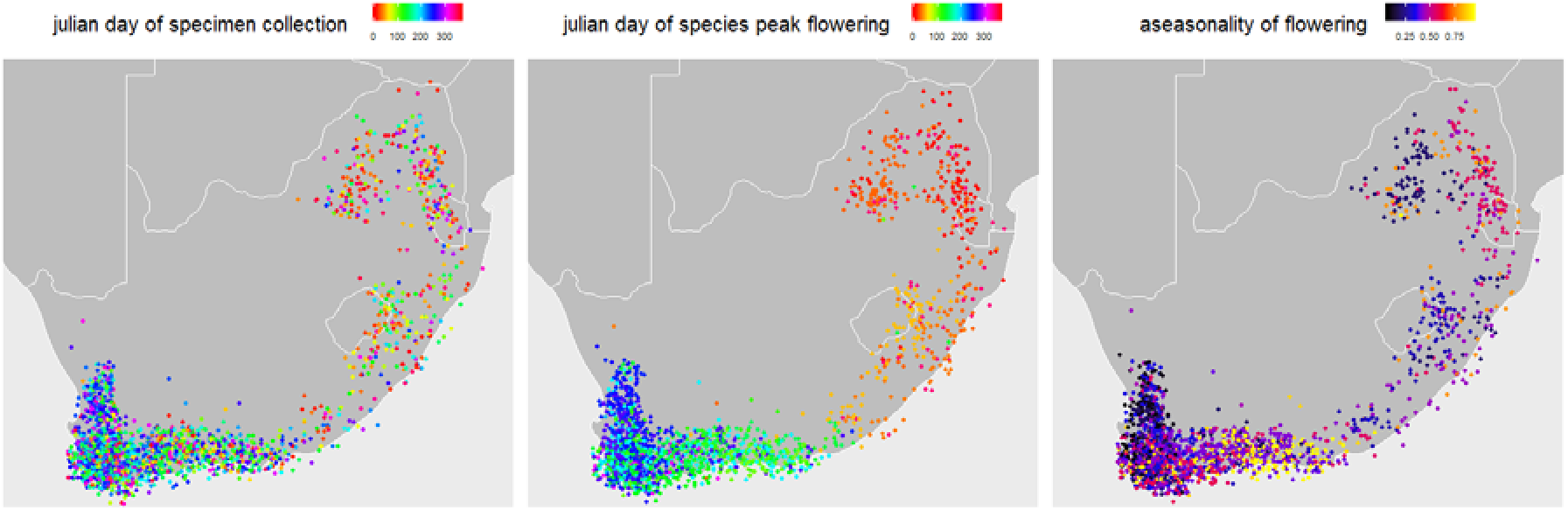
Spatial variability of flowering times of *Protea* species in South Africa derived from preserved herbarium specimens. (A) Day of specimen collection represented in Julian day of year [DOY; where Jan 1 = 1 DOY and Feb 1 = 32 DOY, and so on]. (B) Day of peak flowering, normalized to allow DOY to be treated as a linear variable with no disjunction between Dec 31 and Jan 1, and (C) Aseasonality of flowering to account for species with year-round flowering.

### Effects of climate variation on flowering phenology

For temperature, model selection based on likelihood ratio tests identified highly to marginally significant effects of both spatial variation (p = 0.013) and inter-annual variation (p = 0.058) on specimen collection dates. Neither spatial nor temporal variability in precipitation had significant effects (p = 0.93 and p = 0.75, respectively). The fixed effect coefficients were -3.83 days/°C for spatial climate variation and -5.18 days/°C for year-to-year climate variation, indicating that both dimensions of climatic variability similarly cue early flowering in *Protea* (Figure 4).

**Figure 4.**
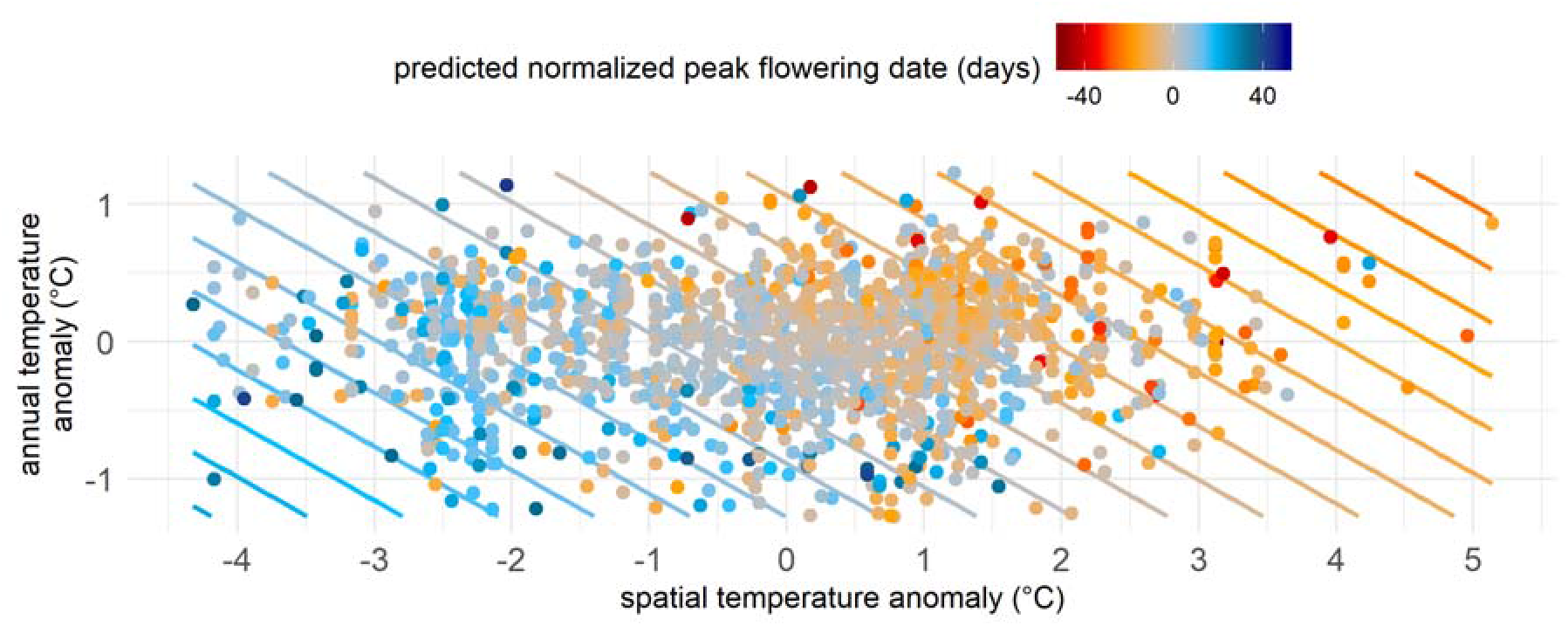
Fitted model predictions of phenological responsiveness of *Protea* species across South Africa as a function of spatial and temporal variation in temperature. Each point represents a *Protea* specimen in the dataset, colored according to its predicted flowering date relative to the species mean. Variability among nearby points is caused by model variables not pictured in the chart, including precipitation variation as well as species-specific responses to temperature. Contour lines represent the overall fixed effects of the two temperature variables on flowering phenology.

In addition to the overall coefficients describing climate effects on phenology, the hierarchical model also includes individual species-level coefficients. Eighty-eight percent of species exhibited advancement in flowering phenology in warmer locations within their ranges while 56% exhibited advancement in warmer years, with well-known species such as *Protea cynaroides* and *P. scolopendrifolia* both showing greatest advancements of -9 days/°C each (Figure S4).

A comparison between a model with only spatial and temporal temperature predictors versus another similar model with an interaction term, showed no significant difference (p = 0.21). Nonetheless, we found a trend toward negative effects of this interaction on flowering phenology, advancing by -2.49 days/°C, hinting that year-to-year temperature variability may have magnified effects in more climatically extreme areas of species ranges.

### Phylogenetic signal in flowering phenology

We tested the hypothesis that closely related species shift flowering time more similarly (either toward early or late flowering) than expected by chance. We found a significant, but weak phylogenetic signal showing that species within lineages shift flowering time more similarly with climate than expected by chance (Figure 5; Abuoheif’s C_mean_ = 0.05, p < 0.01; but both λ = 0.00005 and K = 0.52 [ns]).

**Figure 5.**
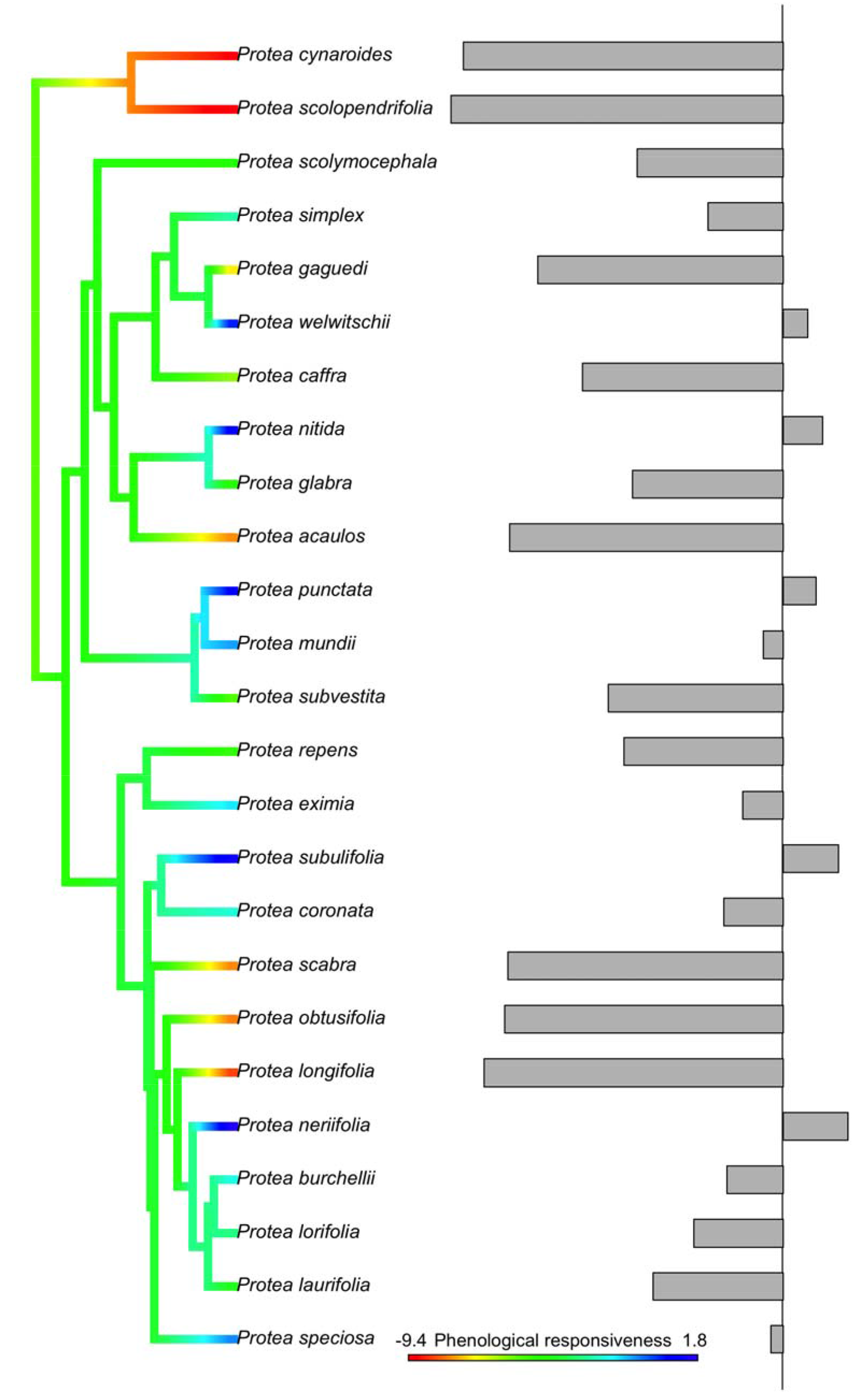
Phylogenetic conservatism of flowering shifts in *Protea*, i.e., the tendency of closely related species to change flowering time similarly under climate change. The color scales correspond to slopes of phenological responsiveness with warming temperature: shifts toward early flowering are indicated in red and late flowering indicated in blue.

## Discussion

In this study, we explored shifts in flowering phenology using herbarium specimens of *Protea* species in southern Africa, providing the first assessment of phenological responses to climate in the southern hemisphere, within an area unrepresented by historic observational data. We show that across temporal and spatial gradients in climate, *Protea* species display advanced flowering phenologies in response to warmer temperature. Although species vary in their phenological responses to climate, these responses are phylogenetically conserved, such that closely related species tended to shift flowering time similarly with temperature.

We found that flowering phenology in *Protea* species advanced by an average of 3–5 days per degree of temperature across both space and time. Our analysis combined these independent dimensions of climate variation—across sites and across years—and found very similar responses of *Protea* species to temperature variation along both dimensions. This strengthens our faith in the results, since simple first principles would indeed predict that these two uncorrelated axes of temperature variation would generate similar flowering phenology responses. It also implies that space-for-time substitution, a widely practiced but less widely validated approach for predicting ecological impacts of future climate change such as species geographic range shifts, may have viability in plant phenology research (Pickett, 1989). Future phenology studies should continue to develop combined independent tests of climatic influence on flowering. During the course of the last 100+ years spanned by our study, South Africa warmed by an average of 1.5 °C (Bomhard et al., 2005; Ziervogel et al., 2014). Such warming temperatures could induce early flowering for the South African *Protea*.

Our results are also in agreement with previous studies from the northern hemisphere showing that plants accelerate their reproductive phenologies with warmer spring temperatures (e.g., Primack et al., 2004; Roberts et al., 2015). These results agree not just qualitatively but are also remarkably similar in effect size, e.g. Primack et al. (2004) and Miller-Rushing and Primack (2008) found slopes of roughly -5 days/°C and -3 days/°C, respectively, for flowering phenology of species in highly seasonal temperate environments of North America. While most such studies have focused on temperate latitudes with seasonal flowering times (Primack et al., 2004; Amano et al., 2010; Panchen et al., 2012; CaraDonna et al., 2014), our study is the first to demonstrate changes in phenology for species with year-round flowering. The congruence between our results for subtropical species with aseasonal flowering to those from temperate regions with highly seasonal phenologies suggests a general trend across disparate ecosystems, and offers an excellent opportunity to explore in future studies of subtropical and tropical ecosystems.

The data preprocessing methods employed here allow for linear regression methods previously used in phenological studies in temperate regions (e.g. Primack et al., 2004, and Miller-Rushing and Primack, 2008) to be applied to more aseasonal systems by using sliding windows to re-center each species’ observations on periods of maximum and minimum flowering activity. When comparing our sliding window assessment of flowering phenology to expert-derived estimates in the literature (Rebelo, 2001), we found very high correlations (r = 0.93), indicating that herbarium records combined with a sliding window approach can indeed capture key phenological patterns. These approaches may thus be of broader use across tropical and subtropical regions, which are often somewhat aseasonal and where most plant diversity occurs, but remain largely unexplored with regard to climate change and phenology.

Phylogenies can provide important insights on species phenological responses to climate change (Davis et al., 2010; Davies et al., 2013). As species are not independent or equivalent evolutionary units due to their shared ancestry, an understanding of phylogenetic patterns of flowering phenology can provide insights into effects of climate change on biological communities. We found that the variability of flowering phenology across species ranges are phylogenetically patterned, such that close relatives have shared phenological responses. While few studies have investigated phylogenetic conservatism in plant phenology, those that have generally find that phenological responsiveness is phylogenetically conserved (Davis et al., 2010; Jia et al., 2011; Lessard-Therrien et al., 2013; Davies et al., 2013). For example, Willis et al. (2008) found that phenological responses to climate among plant clades in New England is phylogenetically conserved such that species within less responsive lineages correspond to those in severe decline. Our results can provide important baselines for more focused investigations of, for example, mechanisms underlying phenological response to climate, and the formation of reproductive barriers that lead to reproductive isolation and sympatric speciation.

In this study, we show that warming cues early flowering in *Protea*, but how this phenomenon influences pollinator abundance remains poorly understood. Members in the iconic genus *Protea* exhibit remarkable variety of floral morphology (Figure 1), most of which have been hypothesized to be maintained by bird and rodent pollinators (Wiens et al., 1983). However, recent studies have also revealed a link for pollination by arthropods (Roets et al., 2006; Steenhuisen and Johnson, 2012). If phenological responsiveness to climate occurs independently among species, some aspects of plant-animal associations including that of pollinators may be modified, potentially leading to phenological mismatch (Kudo and Ida, 2013). We found weak but significant phylogenetic signal in the affinity of closely related species to shift phenology similarly which can potentially influence co-evolved mutualists including principal pollinators. Although we did not explicitly test for phenological responsiveness to warming and pollinator abundance, we believe that herbarium specimens can serve as a critical first step in monitoring species’ population dynamics including distributional migrations, expansions, and contractions (Feeley, 2012), but also as a roadmap to explore pollinator phenological responsiveness to climate change.

The use of herbarium records to investigate phenological response with climate adds a new perspective to phenological investigations in areas unrepresented by historic observational data such as the southern hemisphere. Observational data have informed much of our understanding of biological responses to climate change. However, observations-only records are seldom validated by specialists as opposed to vouchered herbarium specimens, which offer a means of taxonomic validation by specialists, making them important for the integrity of biodiversity analyses. Historical field observations are limited, whereas herbarium data capture dynamics of the past and present and can be used to predict the future. Despite various weaknesses and biases associated with the sampling of herbarium specimen (Daru et al., 2018; Meineke et al., 2018a,b), they are an untapped resource for understanding changes in phenology with climate. As such they can be used as a baseline pending when real-time monitoring data becomes available. Finally, our analyses reveal the existence of previously unidentified patterns of phylogenetically non-random flowering phenology, reinforcing the need to study phenology in a phylogenetic context.

## Acknowledgements

We thank Thandy Makgolane for help with data collation. We are particularly grateful to the National Herbarium of the South African National Biodiversity Institute (http://newposa.sanbi.org/) for granting us access to their data. B.H.D thanks Texas A&M University-Corpus Christi for start-up funds and logistic support.

## Supplementary Information

Figures S1-S7 are available in the following pages

**Figure S1.**
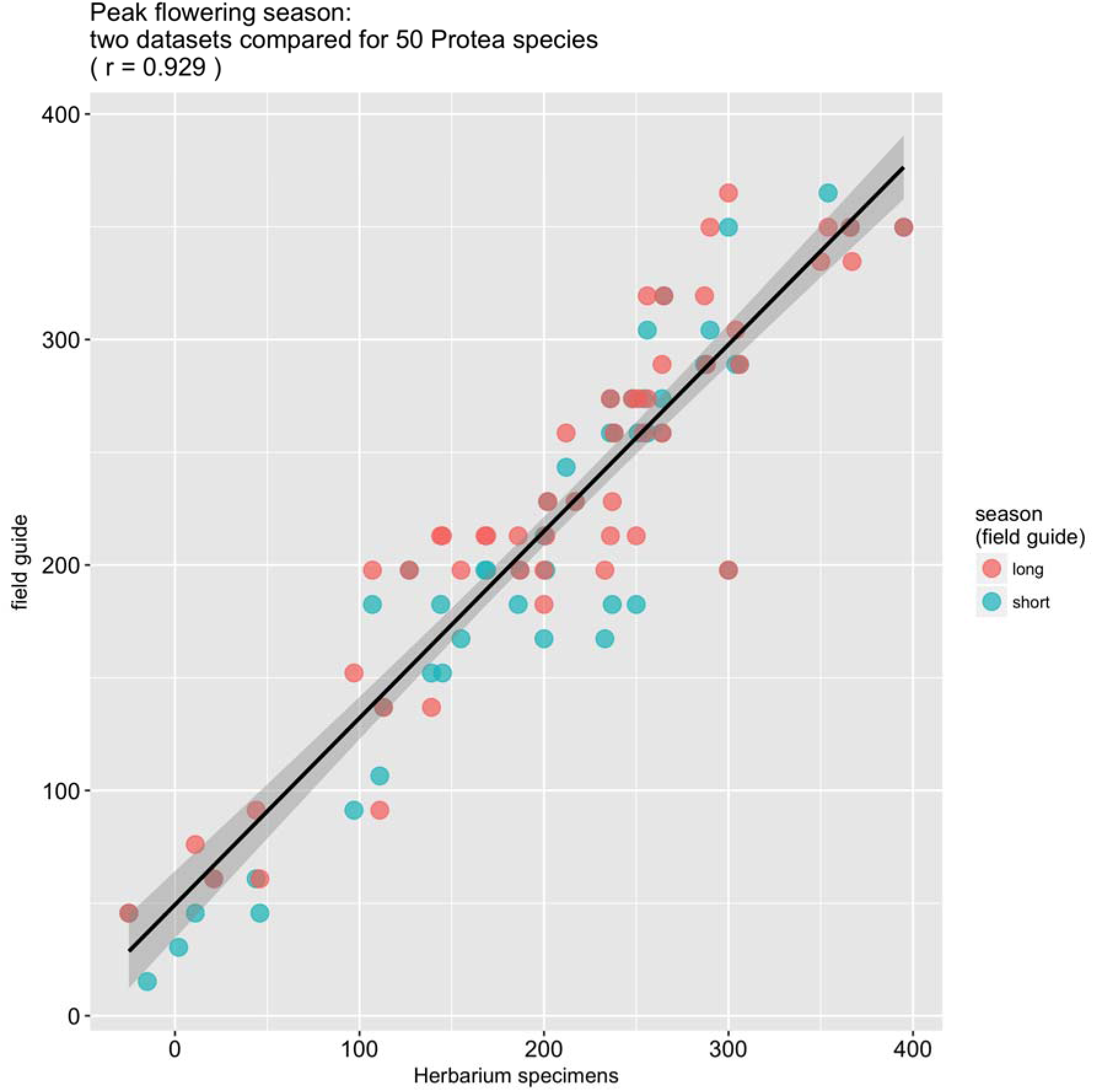
Comparison of species peak flowering season (in Julian days) recorded from herbarium specimen records versus literature (Rebelo 2001). Rebelo (2001) reports both a long and short flowering season for each species, the centers of which are shown here relative to the peak flowering date we calculated from herbarium data as described in the text. While the y-axis ranges from 0–365, the x-axis has a slightly broader range---given the circular nature of the calendar year, a given Julian date can take multiple values (e.g. 10 = 375), and the value that best communicates alignment with the field guide dataset is shown.

**Figure S2.**
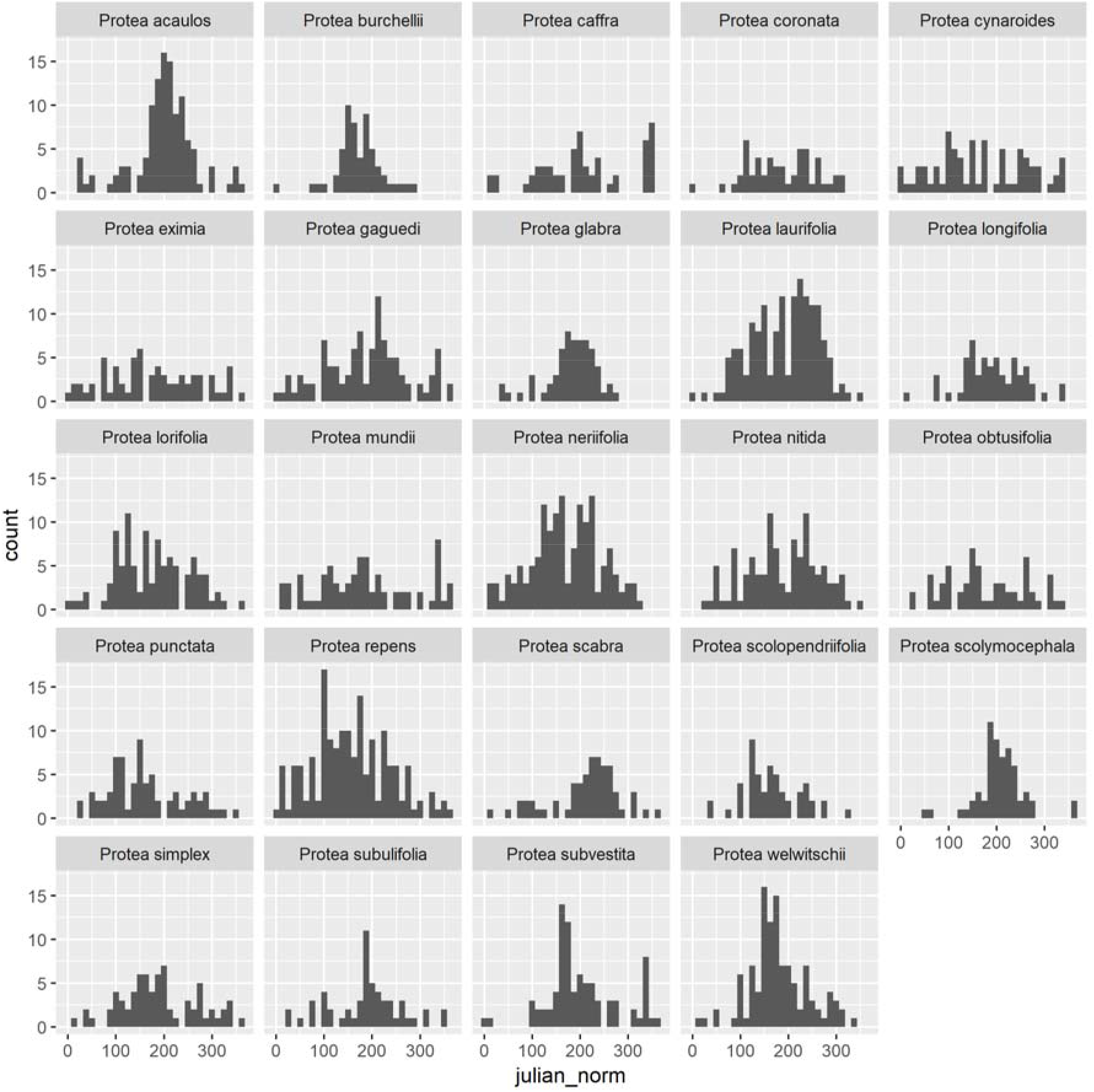
Specimen collection frequency across “day of flowering year”, a normalized version of the Julian day of year.

**Figure S3.**
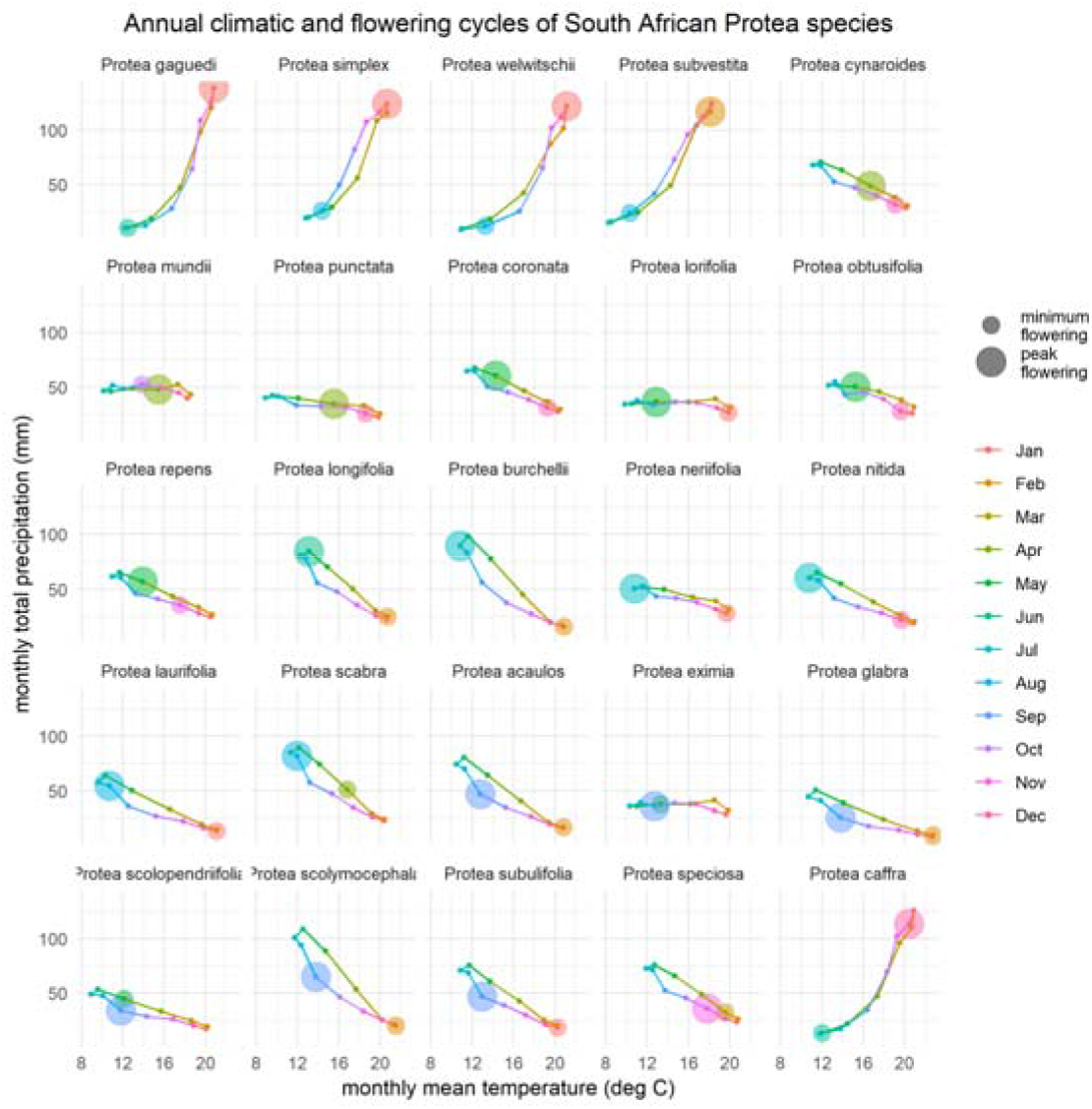
Monthly mean temperature and precipitation and flowering phenology of the 25 Protea species included in the analysis.

**Figure S4.**
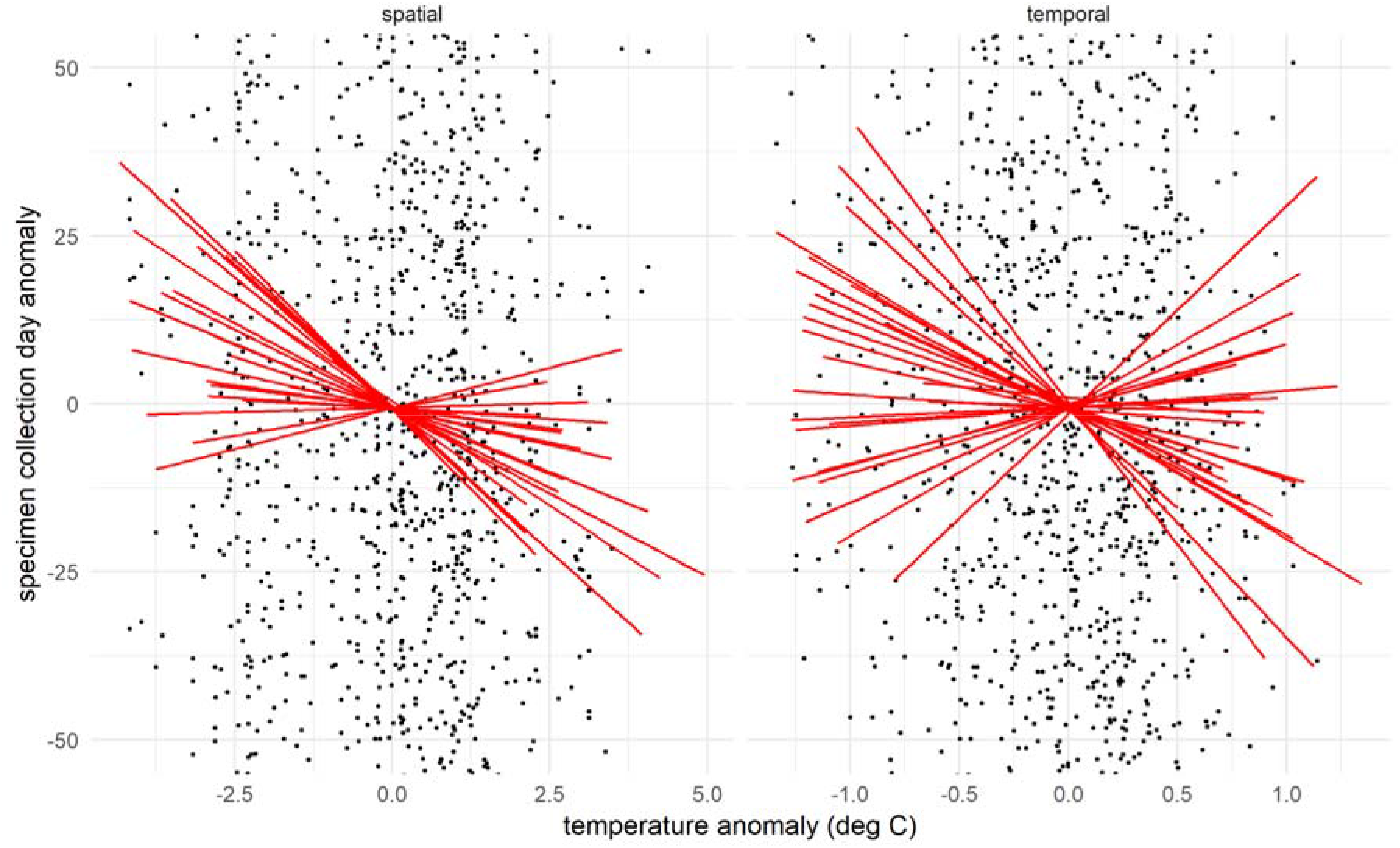
Changes in flowering times of *Protea* species across South Africa in relation to anomalies in temperature. Statistical analysis based on mixed effects model using both (A) spatial temperature variation, and (B) temporal climate (year-to-year temperature variation) as predictors, with species as random effect. Negative slopes indicate advancement of flowering with warming. Lines indicate fitted slopes for individual Protea species. Points indicate input specimen data, and have been truncated for visualization at the extremes of the y-axis range.

